# The Use of Visual Cues in Gravity Judgements on Parabolic Motion

**DOI:** 10.1101/301077

**Authors:** Björn Jörges, Lena Slupinski, Joan López-Moliner

**Author notes:** Correspondence, **Passeig de la Vall** d’Hebron, 171 08035 **Barcelona**.

## Abstract

Evidence suggests that humans rely on an earth gravity prior for sensory-motor tasks like catching or reaching. Even under earth-discrepant conditions, this prior biases perception and action towards assuming a gravitational downwards acceleration of 9.81 m/s^2^. This can be particularly detrimental in interactions with virtual environments employing earth-discrepant gravity conditions for their visual presentation. The present study thus investigates how well humans discriminate visually presented gravities and which cues they use to extract gravity from the visual scene. To this end, we employed a Two-Interval Forced-Choice Design. In Experiment 1, participants had to judge which of two presented parabolas had the higher underlying gravity. We used two initial vertical velocities, two horizontal velocities and a constant target size. Experiment 2 added a manipulation of the reliability of the target size. Experiment 1 shows that participants have generally high discrimination thresholds for visually presented gravities, with weber fractions of 13 to beyond 30 %. We identified the rate of change of the elevation angle (*ẏ*) and the visual angle (*θ*) as major cues. Experiment 2 suggests furthermore that size variability has a small influence on discrimination thresholds, while at the same time larger size variability increases reliance on *ẏ* and decreases reliance on *θ*. All in all, even though we use all available information, humans display low precision when extracting the governing gravity from a visual scene, which might further impact our capabilities of adapting to earth-discrepant gravity conditions with visual information alone.

## Introduction

Improvements in applicability and cost-efficiency of Virtual and Augmented Reality technologies have led to a surge in their popularity. More and more applications are pushing boundaries by immersing users into worlds that defy the regularities of our natural environment. Among these pervasive laws is the pull of gravity, which is ubiquitous and almost invariant across the world (9.78 m/s^2^ at the Equator and 9.832 m/s^2^ at the poles). Our life-long exposure to this specific value gives rise to the concern that altered gravity values might pose a significant challenge to users. And in fact, perceptuo-motor performance under earth-discrepant gravity conditions has been receiving some attention over the past decades: Prominently, an internal representation of earth gravity has been suggested to be involved in a series of sensory-motor tasks such as catching and reaching. While arbitrary accelerations are generally not picked up by the perceptual system (Brenner et al., 2016; Werkhoven, Snippe, & Alexander, 1992), humans can make use of this gravity prior to improve catching performance for objects accelerated by earth gravity. This discrepancy between arbitrary accelerations and acceleration through earth gravity is particularly salient when online information is not available (partially occluded trajectories) or unreliable (noisy presentation). The utility of such model has been substantiated in numerous ways: (McIntyre, Zago, Berthoz, & Lacquaniti, 2001; McIntyre, Zago, Berthoz, & Lacquaniti, 2003) showed that even after extensive exposure to zero gravity in space, catching movements were initiated too early with regard to Time-to-Contact for balls dropping at a constant speed, indicating that humans rely on their representation of earth gravity even when visual and bodily cues indicate a discrepant gravity. A series of studies conducted in a semi-virtual task on earth (Zago et al., 2004; Zago & Lacquaniti, 2005) demonstrated that even after extensive training over up to two sessions, participants did not fully adapt to visually presented zero gravity and were still expecting targets to accelerate downwards. Even remembered locations of horizontally moving projectiles seem to drift downwards, in direction of earth-gravity, over time (De Sá Teixeira, Hecht, & Oliveira, 2013). Also, brain imaging and lesion studies have showed areas differentially activated for (Indovina et al., 2005) or dedicated to (Maffei et al., 2016) computations involving earth-gravity. While concerns have been raised about the parsimony of this way of framing the results (Baurès, Benguigui, Amorim, & Siegler, 2007; but see also (Zago, McIntyre, Senot, & Lacquaniti, 2008) for a rebuttal), the overall picture remains intact: There is evidence that an internal representation of earth-gravity is accessed and applied even when this is to the detriment of the performer. While this internal model of earth gravity has been studied thoroughly, it remains largely unknown how humans (would) extract the underlying gravity value from the dynamics of a visual scene. To bridge this gap in our understanding, the present study takes a look at how well the visual system computes gravity from observing its effects on the objects in a virtual environment. Furthermore, we scrutinize the roles of different visual and temporal cues humans may rely upon for their decision.

On a more theoretical level, our study aims at interpreting gravity perception judgements within a Bayesian framework. According to this framework, sensory information (“likelihood”) is integrated with previous knowledge about the world (“prior”), yielding a more precise and usually more accurate final percept (“posterior”). The weights of likelihood and prior are a function of their respective reliability. Within this framework, the internal model of gravity can be described as a so called strong prior (Jörges & López-Moliner, 2017): as the evolution of the human species as well as the development of every single human took place under a largely invariant gravity value of 9.81 m/s^2^, the reliability of this prior is extremely high. It thus overrules all sensory information represented as the likelihood. However, the experimental results cited above only imply a strong relative weight of the prior with regards to the likelihood; this is also consistent with a weak likelihood combined with an average prior or a weak likelihood combined with a strong prior. While some evidence has been provided that visual acceleration information is relatively unreliable (Benguigui, Ripoll, & Broderick, 2003; Brenner et al., 2016; Werkhoven et al., 1992), the nature of the likelihood remains to be investigated specifically for gravitational accelerations.

## Experiment 1

### Materials and Methods

#### Participants

A total of eleven (n = 11) participants performed the task, among them two of the authors (BJ and JLM). All had normal or corrected-to-normal vision. One (n = 1) subject was excluded because they didn’t follow instructions and another (n = 1) was excluded because their performance was at chance level for all stimulus strengths and a post-hoc stereo-vision test revealed that they were stereo-blind. The remaining participants were in an age range of 19 and 51 years and five (n = 5) were female. We did not test their explicit knowledge of physics, as previous studies suggest that explicit knowledge about gravity has no effect on performance in related tasks (Flavell, 2014; Kozhevnikov & Hegarty, 2001). All participants gave their informed consent. The research in this study is part of an ongoing research program that has been approved by the local ethics committee of the University of Barcelona. The experiment was conducted in accordance with the Code of Ethics of the World Medical Association (Declaration of Helsinki).

#### Apparatus

Two Sony laser projectors (VPL-FHZ57) were used to provide overlaid images in a back-projection screen (244 cm height and 184 cm width) with a resolution of 1920×1080 pixels. The frequency of refresh of the image was 85 Hz for each eye. Circular polarizing filters were used to provide stereoscopic images. Participants stood at 2 m distance centrally in front of the screen and were using polarized glasses to perceive the object stereoscopically. The shown disparity was adapted to each participant’s inter-ocular distance.

#### Stimuli

Our stimuli were spheres of tennis ball size (r = 0.033 m) that approached the observers frontally on parabolic trajectories. They could be governed by one of seven test gravities between 0.7 and 1.3g (in steps of 0.1g). Furthermore, they could have two different initial vertical velocities (3.7 and 5.2 m/s) and two different horizontal velocities (6 and 8.33 m/s); see Figure 1. As air drag was neglected, the horizontal velocity remained constant along the trajectory, while the vertical velocity changed according to the governing gravity. The starting z position varied in function of horizontal velocity and gravity, such that the endpoint was always the observer. The starting position and endpoint in the y dimension was 56 cm above the ground. As 3D presentation in our setup breaks down when the projected target gets too close to the observer, the ball disappeared at a random point within the last 88 to 98 % of the trajectory. This corresponds to 88 to 98 % of the overall time it would take for the target to reach the observer. Each parabola’s presentation time was determined by the time-to-arrival, which in turn depended on initial vertical velocity and gravity. We did not include a detailed visual scene because we were focusing on the visual parameters (elevation angle, visual angle and their temporal derivatives) changing with different gravities. A detailed 3D visual scene may have shifted the processing focus away from visual parameters and towards physical values such as the overall distance or height, as they can be recovered more easily when a stimulus is presented in a rich environment.

**Figure 1:**
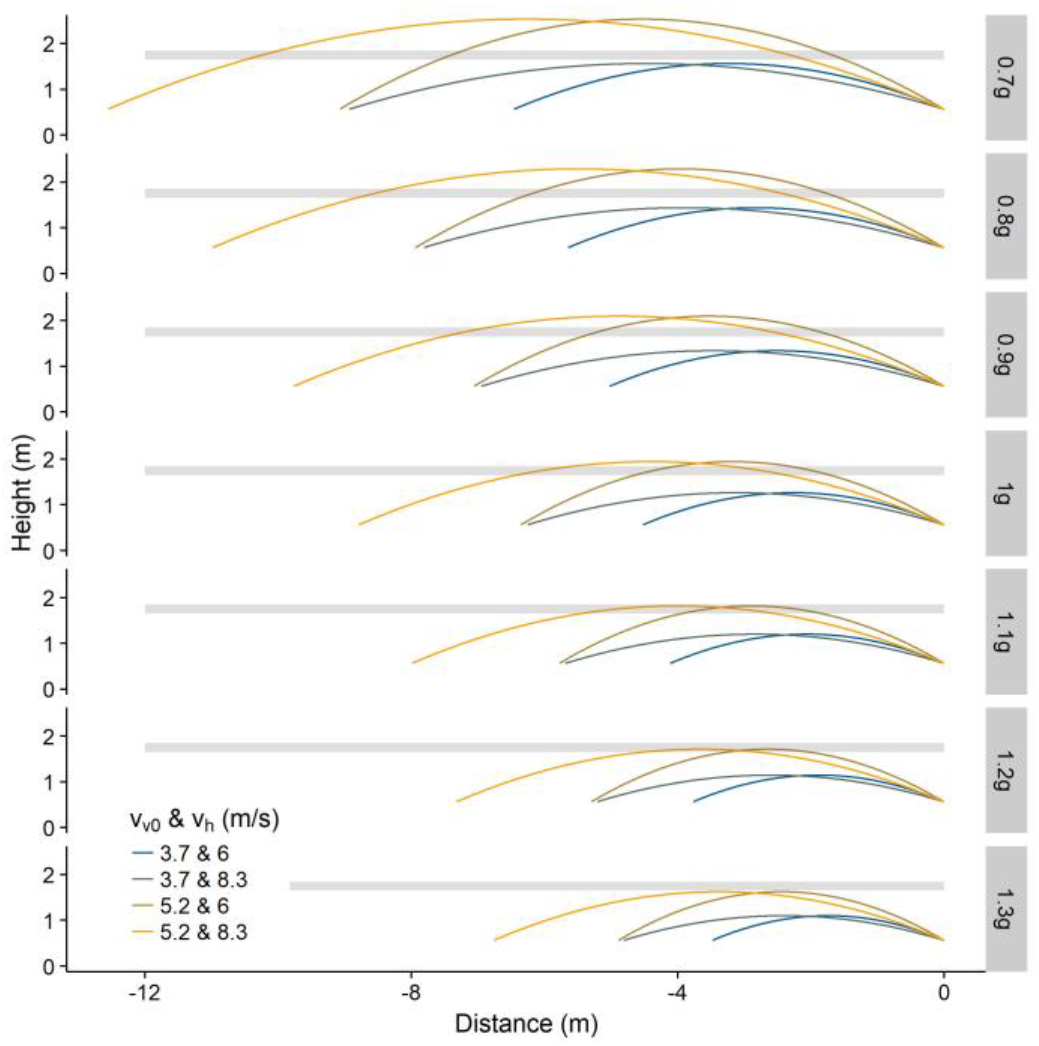
Lateral view of the spatial trajectories of the parabolas that served as stimuli. The panels represent different gravity values ranging from 0.7g to 1.3g in steps of 0.1g. Spatial differences are due to different initial vertical velocities (3.7 or 5.2 m/s) and different horizontal velocities (6 or 8.33 m/s). The shaded rectangles designate the range of eye-levels of participants (1.65 m – 1.88 m). The observer’s position is a x = 0 m.

#### Procedure

We employed a Two-Interval Forced-Choice (2IFC) task. Participants were asked which of two presented parabolas had the greater underlying gravity. Each trial consisted of two parabolas in random order: One standard (or reference) parabola (1 g) and one test parabola (one out of the seven gravity test values). The order of presentation of the two parabolas was randomized, but the method guaranteed that each of the seven gravity test values was presented the same number of times (28 times per block, 112 times in total). Initial vertical and horizontal velocities were allotted to the parabolas randomly and independently of each other. After each trial, the participants heard a beep that indicated they could give their response by pressing one of two mouse buttons (left: first parabola had greater underlying gravity; right: otherwise). The click response also initiated the next trial. 560 trials were presented in four blocks of 140 trials. Before starting the actual experiment, participants were given instructions and did up to 15 familiarization trials. Where necessary, we explained the concept of gravity both theoretically and with real life examples. We only started the main body of the experiment when participants confirmed that they were comfortable enough with their understanding of gravity. At no point, feedback on the accuracy was provided.

#### Optic flow analysis

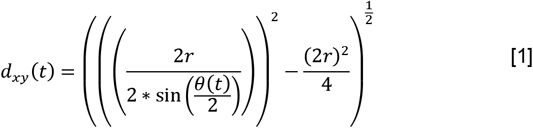

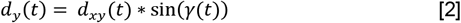

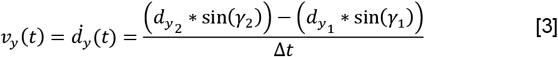

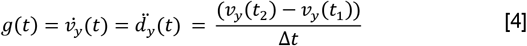

We furthermore provide an evaluation of the geometry of parabolic trajectories in order to identify optic variables that could be used to decode underlying gravities. We identified two ways to recover the gravity value from visual and temporal information. The first one relies on successive sampling of visual angle and elevation angle information: (Equation 1) shows how the distance (within the xy-plane) to a round object (*d_xy_*) can be recovered unambiguously from the visual angle (*θ*) when its size (r = radius) is known. (Equation 2) shows how the y position (*d_y_*) with respect to eye level can be recovered from the Elevation Angle (*γ*) and the distance to the ball (*d_xy_*). Finally, (Equation 3) and (Equation 4) demonstrate how second-order information about the downwards acceleration of the object can be extracted through successive sampling of the y position. It is thus possible to recover gravity from combining visual angle and elevation angle information over time when a stable representation of the target size is maintained. Note, however, that subjects do not only have to recover the vertical velocity from the optical cues, but also estimate its temporal derivative. It has been shown before that participants quickly establish an accurate representation of the size of virtual objects they deal with (Hosking & Crassini, 2010; López-Moliner, Field, & Wann, 2007; López-Moliner & Keil, 2012). This is facilitated when stereoscopic information is available (as in our case; Regan & Beverley, 1979). We can thus assume that our participants maintained a relatively accurate representation of the ball size. The known size can then be used in combination with the visual angle to recover the distance accurately. While thus all information necessary for distance recovery is readily available to the subject, it has been observed that distance may be underestimated in virtual environments, especially when these are scarce (Loomis & Knapp, 2003; Messing & Durgin, 2005). While the visual angle itself can be estimated with accuracy (McKee & Welch, 1992), its rate of change over time is very noisy, with Weber fractions of 10 % and beyond (Gómez & López-Moliner, 2013; Regan & Hamstra, 1993). Nonetheless, as suggested by the important, though contested role of the visual angle and its rate of expansion (for the parameter τ) in the literature on TTC estimation for linearly approaching targets (Lee & Reddish, 1981), it may still deliver some information, e. g. about the physical horizontal velocity. The elevation angle in turn is readily picked up by the visual system as long as the target is clearly visible. Note that we define the elevation angle as angle between a line through the observer’s eyes and the starting position on the one hand, and a line through the observer’s eyes and the target at time t on the other hand. An alternative possibility would be to use a fixed reference in the world (such as the observer’s straight ahead on eye-level), but in our opinion it is more likely that participants use the starting point as reference for the rest of the parabola. Furthermore, its rate of change directly corresponds to retinal speed, which makes the recovery of this optic variable relatively easy. Previous research on human sensitivity to the rate of change of the elevation angle shows that it can be detected with a 5 % error margin when it is between 0.03 and 1.2 rad/s (McKee, 1981). Figure 2A illustrates that 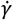 has a zero-crossing between 25 % and 50 % of the trajectory, depending on the velocity profile, which indicates that, in the first half of each parabola, there is a part where 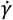 is smaller than 0.03 rad/s and thus very noisy. Furthermore, (the absolute value of) 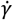 grows beyond 1.2 rad/s after 50 % of the trajectory, which again makes estimates more noisy, with reported Weber fractions of 12 % for 2.23 rad/s and 22 % for 4.46 rad/s (de Bruyn & Orban, 1988). However, differences between the gravities become much more pronounced after this point, which suggests that recovering the gravity value is facilitated in the second part of the trajectory. It is important to add that, as we did not record eye-movements, we use the physical values of the optic parameters as approximations of the parameters actually perceived and represented by the participants. Note also that we calculated the optic values for each participant individually according to their eye-height (see also the shaded rectangle in Figure 1, denoting the range of eye-heights).

**Figure 2:**
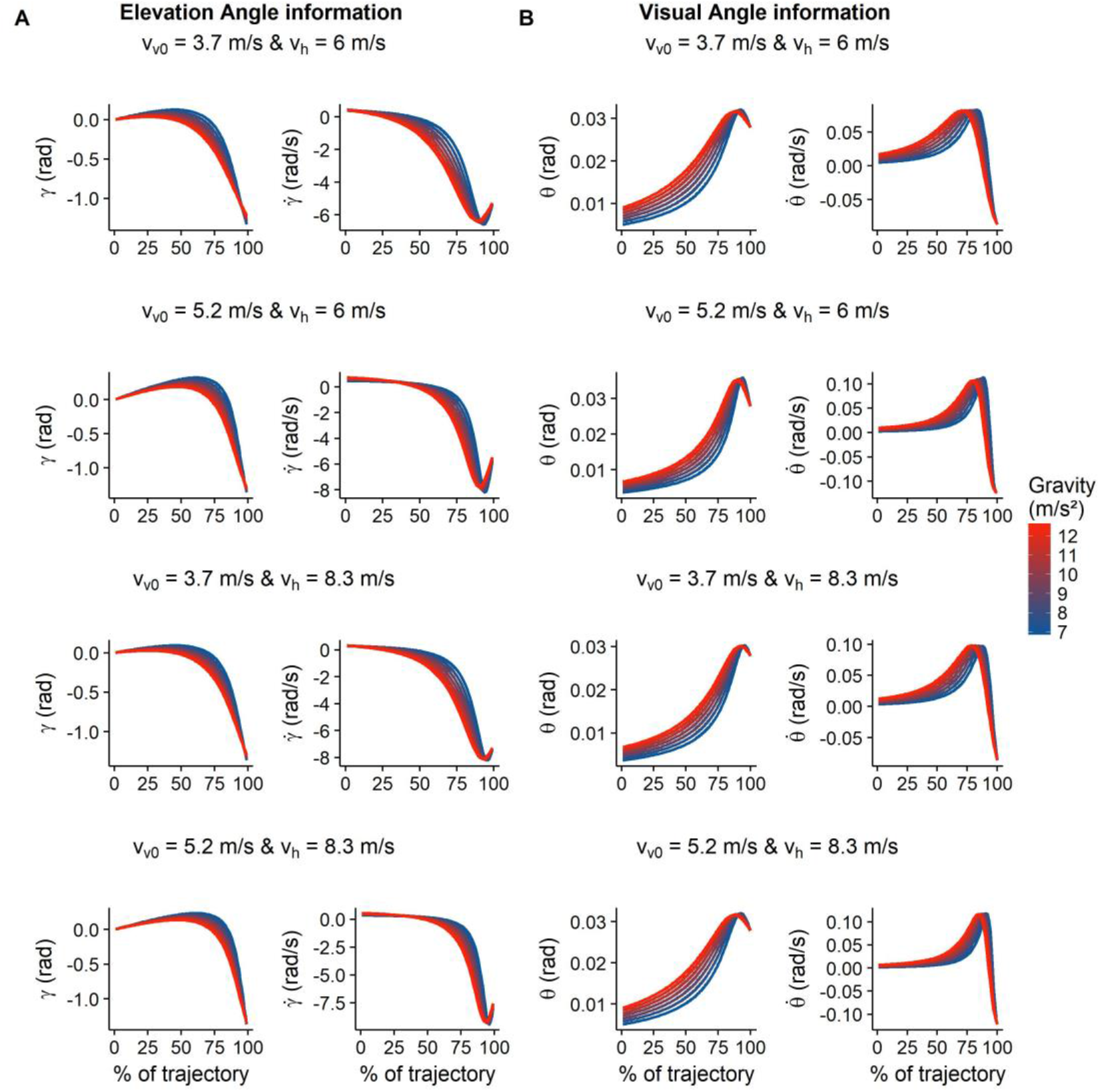
A. The time course of Elevation Angle information and its derivative (*γ* and 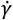) for different gravities (color gradient), plotted in different panels for each velocity profile used in our experiment. B. Development of Visual Angle information (*θ* and 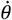) over the course of the trajectory for different gravities (color gradient), plotted in different panels for each velocity profile used in our experiment.

#### Data analysis

We fitted psychometric functions to our data. We used the gravity test values as well as differences between test and standard parabola in elevation angle (*γ*), the temporal derivative of the elevation angle 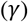, the visual angle (*θ*) and the temporal derivative of the visual angle 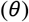 as decision variables to model responses. Decision variables are optical cues that participants can use to make judgments. While *γ*, 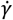, *θ* and 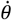 may be picked up directly by the visual system, gravity needs to be estimated by integrating different optic and temporal cues. When talking about the decision variable Gravity, we therefore refer to it as a placeholder for an approximately accurate combination of the available cues (see Equations (1) – (4)), as opposed to subjects relying solely or predominantly on one optic flow cue.

We used the R (R Core Team, 2017) package QuickPsy (Linares & López-Moliner, 2016) to fit cumulative Gaussians. We maximized the log-likelihood to obtain the best set of parameters for the mean of the cumulative Gaussian, which can be used to assess bias or the Point of Subjective Equality (PSE), and its standard deviation, which provides a means to assess discrimination thresholds. When a standardized scale is used for the decision variables, the standard deviations can also be used to assess how well a decision variable explains variability in participant responses in comparison to other decision variables. Where relevant, we therefore standardized stimulus values, dividing them by the stimulus range of the respective decision variable. We furthermore assessed how well the model fit the data by means of the Akaike Information Criterion (AIC), which serves as another indicator of how much participants’ judgments relied to the respective cue.

We first report the overall sensitivity for gravity judgements and analyze whether the different initial velocity profiles lead to biases or differences in precision. Then, using model fits, we determine a region of interest throughout the parabolas on which participants base their judgments principally, followed by an analysis of which variant of the optic flow parameters *γ* and *θ* is most useful to predict performance. The next step is an analysis on the subject level for the chosen cues. Then, we try out different combinations of *γ* and *θ*, which, according to a gravity-based TTC model (“GS model”; Gómez & López-Moliner, 2013), might be used as approximate heuristics to estimate gravity. Finally, we employ Generalized Linear Mixed Modelling (Moscatelli & Lacquaniti, 2012) to assess whether participants used homogeneous or rather idiosyncratic strategies, and finish with a short assessment of the role of purely temporal information.

### Results

#### Gravity judgements and physical velocities

Fitting psychometric functions across data of all subjects and velocity conditions and with gravity as decision variable, we obtained a PSE of 9.91 m/s^2^ (95 % CI = [9.80, 10.03]) and a standard deviation of 3.45 m/s^2^ (95 % CI = [3.23, 3.66]; corresponding to a Weber fraction of 23.8 %). As first step deeper into the analysis, we compared PSEs and discrimination thresholds between different initial velocity profiles. To this end, we split the data by differences in initial vertical velocity between test and standard parabola (Figures 3A, B, C), and by differences in horizontal velocity between test and standard parabola (Figures 3D, E, F). Figure 3A shows the probability to judge the test parabola’s gravity as larger than the standard parabola’s gravity, averaged across subjects and split by differences in initial vertical velocities. Figure 3D shows the same data but split by differences in horizontal velocities. Figure 3B and 3C display the PSE (3B) and standard deviations (3C) of the psychometric curves depicted in 3A, while Figure 3E and 3F depict PSE (3E) and standard deviations (3F) for the psychometric curves depicted in 3F.

**Figure 3:**
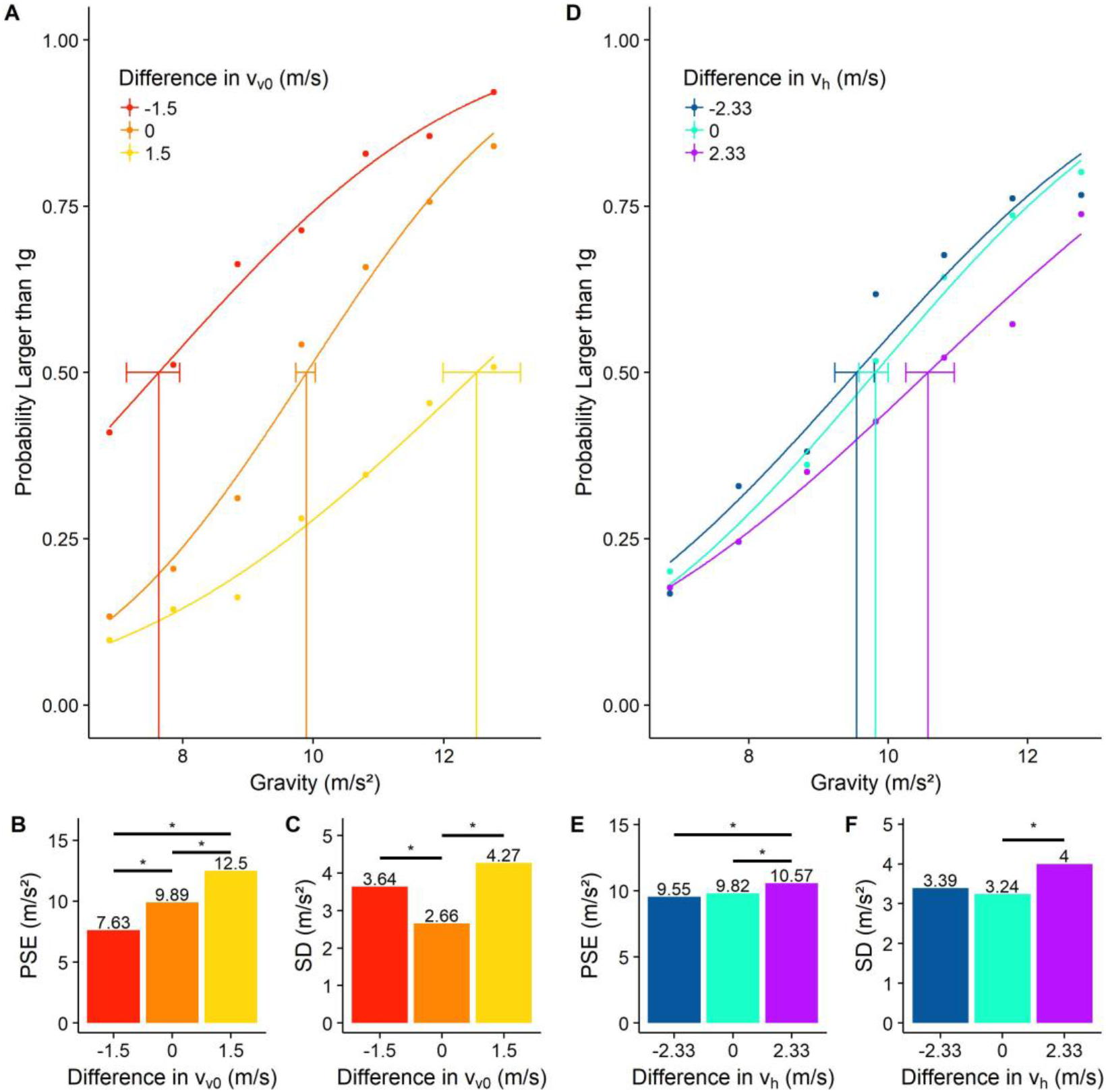
**A, B, C**: Psychometric functions for data split by difference in vertical velocity between test and standard parabola (A), PSEs (B) and SDs (C). **D, E, F**: Psychometric functions for data split by difference in horizontal velocity between test and standard parabola (A), PSEs (B) and SDs (C). For A and D, the stimulus strength is plotted against the probability to judge the test parabola as having the higher underlying gravity. Horizontal bars in A and D indicate the confidence interval for the PSEs. Horizontal bars and stars in B, C, E and F indicate that the 98.3 % confidence intervals around the respective difference estimates did not contain zero (see text for a description of the statistics).

We employed the Bootstrap (Efron & Tibshirani, 1998) implemented in QuickPsy in order to test for biases and differences in discrimination thresholds between the different conditions. This method calculates confidence intervals for estimations of PSE and SD differences between all conditions, based on a significance level. If a confidence interval doesn’t contain zero, there is a significant difference between the groups. We corrected for multiple comparisons with the Bonferroni method (Bland & Altman, 1995), that is, we raised the significance level of the confidence intervals to 1-0.05/n, with n being the number of comparisons.

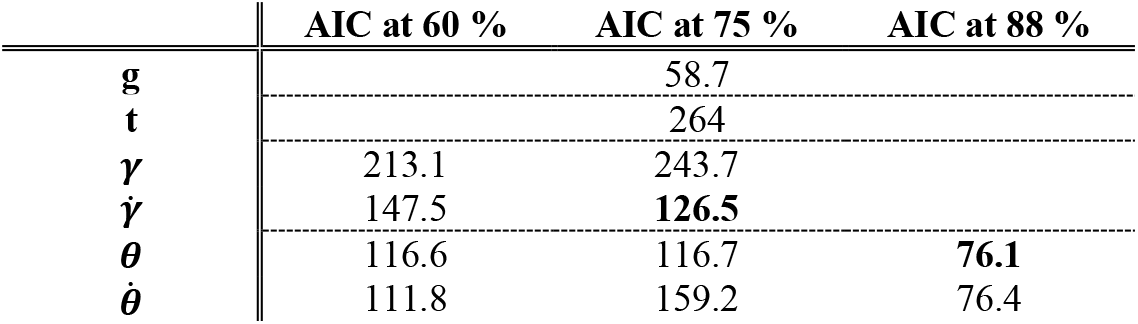

|  | AIC at 60 % | AIC at 75 % | AIC at 88 % |
| --- | --- | --- | --- |
| **g** |  | 58.7 |  |
| **t** |  | 264 |  |
| **$\gamma$** | 213.1 | 243.7 |  |
| **$\dot{\gamma}$** | 147.5 | **126.5** |  |
| **$\theta$** | 116.6 | 116.7 | **76.1** |
| **$\dot{\theta}$** | 111.8 | 159.2 | 76.4 |

Initial vertical velocities had a striking influence both on PSEs and on SDs: PSEs differed significantly between all three conditions (*PSE*_−1.5_ − *PSE*_0_ = −2.27 m/s^2^, CI = [−2.83,−1.92]; *PSE*_−1.5_ − *PSE*_1.5_ = 4.88 m/s^2^, CI = [−5.70,−4.21]; *PSE*_0_ − *PSE*_1.5_ = −2.61 m/s^2^, CI = [−3.30,−2.04]; all confidence intervals based on a significance level of 1-0.05/3 = 0.983), illustrating a strong bias to judge parabolas with higher initial vertical velocities as having the lower gravity. The SDs differed significantly between two condition pairs (*SD*_−1.5_ − *SD*_0_ = 0.98 m/s^2^, CI = [0.46,1.78]; *SD*_0_ − *SD*_1.5_ = 1.61 m/s^2^, CI = [-2.60,−0.77]; all confidence intervals based on a significance level of 1-0.05/3 = 0.983), indicating that discrimination thresholds were lower when the initial vertical velocities of test and standard parabola were the same.

Also horizontal velocities impacted performance, but more weakly than vertical velocities. PSEs were significantly different for two comparisons (*PSE*_−2.33_ − *PSE*_2.33_ = −1.02 m/s^2^, CI = [−1.43,−0.57]; *PSE*_0_ − *PSE*_2.33_ = −0.75 m/s^2^, CI = [−1.06,−0.30]; all confidence intervals based on a significance level of 1-0.05/3 = 0.983), indicating a bias to judge parabolas with higher horizontal velocities as having the higher gravity. Regarding the SDs, only one comparison turned out significant (*SD*_0_ − *SD*_2.33_ = −0.77 m/s^2^, CI = [−1.62,−0.09]; confidence interval based on a significance level of 1-0.05/3 = 0.983), illustrating lower discrimination thresholds when both horizontal velocities coincided than when the test parabola’s horizontal velocity was higher than for the standard parabola.

These results show that initial vertical velocities have a substantial impact on performance: higher initial vertical velocities bias participants to judge these parabolas as having the lower underlying gravity and vice-versa. Furthermore, discrimination thresholds decreased when both test and standard parabola had the same initial vertical velocity. The horizontal velocity only had a minor impact.

#### Optic flow cues

We then proceeded to analyze the visual parameters that convey information about the gravity underlying parabolas. As discussed before, we identified the elevation angle and the visual angle and their respective derivatives as possible candidates for analysis. We thus fitted psychometric models for (1) the elevation angle (*γ*), (2) the temporal derivative of the elevation angle 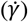, (3) the visual angle (*θ*), and (3) the temporal derivative of the visual angle 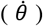. As both sources’ informational content varies throughout the trajectory, we had to choose a specific point of the parabola for which to conduct the analyses. As described above, 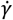 could be estimated with a 5 % error margin throughout the first 60-70 % of the trajectory, as per the 5 % discrimation threshold between 0.03 and 1.2 rad/s reported by McKee (1981). 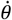 is very small during the first half of the trajectory, which may further impact its estimation; in general, it has been shown to be extremely noisy until it gets very close to the observer (Gómez & López-Moliner, 2013). Estimating *γ* and *θ* is, in turn, not very noisy (McKee & Welch, 1992). Finally, Figure 2 illustrates that gravity-based differences in optical cues become stronger in later parts of the parabolas. Based on these considerations, we fitted models with the difference in *γ*, 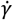, *θ* and 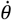 at 60 % and 75 % of the trajectories between test and standard parabola as decision variables, as well as *θ* and 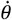 at 88 % of the trajectories, which represents the moment in which visual angle information was most readily available and the targets were still visible for every trajectory.

Among elevation angle cues, 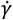 achieved the best model fits at 75 % of the trajectory (see Table 1), while the visual angle cues *θ* and 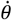 had the best fits at 88 %, with a neglectable difference between both. The cues at 60 % displayed generally worse fits. This analysis confirms that information in the last third of the trajectory is privileged. Despite visual angle fits being superior at 88 % with regards to 75 %, we chose *θ* at 75 % for further analyses, as its fit was still acceptable and the literature suggests that a combination of 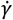 and *θ* may play a privileged role in perceptual processes related to parabolic trajectories (Gómez & López-Moliner, 2013).

We then proceeded to a subject-level analysis for the placeholder decision variable gravity and the basic optical cues 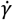 and *θ*. Figure 4 displays the psychometric functions for each subject based on differences in gravity (4A), 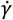 at 75 % of the trajectory (4C) and *θ* at 75 % of the trajectory (4E), as well as the respective SDs (4B, D, F). The PSEs were not included in the figure because they did not differ significantly from the mean value for any of the decision variables, which is in line with the absence of any biasing factors. We standardized the values for this analysis in order to facilitate a comparison between the variability explained by gravity, 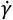 and *θ* respectively. Fitting the data per subject revealed a huge inter-subject variability in discrimination thresholds: SDs ranged between 0.33 m/s^2^ and 1.37 m/s^2^ for gravity. A similarly high inter-subject variability could be observed for 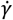, where SDs ranged between 0.26 rad/s and 1.24 rad/s, and for *θ*, where SDs ranged between 0.3 rad and 1.38 rad. Interestingly, SD patterns do not match between the three decision variables: for some subjects (s01, s05), gravity clearly explained more variability than 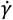 nor *θ* – as evidenced by lower SDs –, suggesting that they combined cues for more accurate judgements. For others (s02, s09, s11), however, particularly 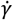 made for much more saturated psychometric functions than gravity, suggesting that these subjects relied mainly on 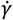 in their judgements. *θ* followed the pattern of 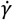 approximately (with the exception of s02) and our results do not allow to establish a clear primacy of one cue of the other.

**Figure 4.**
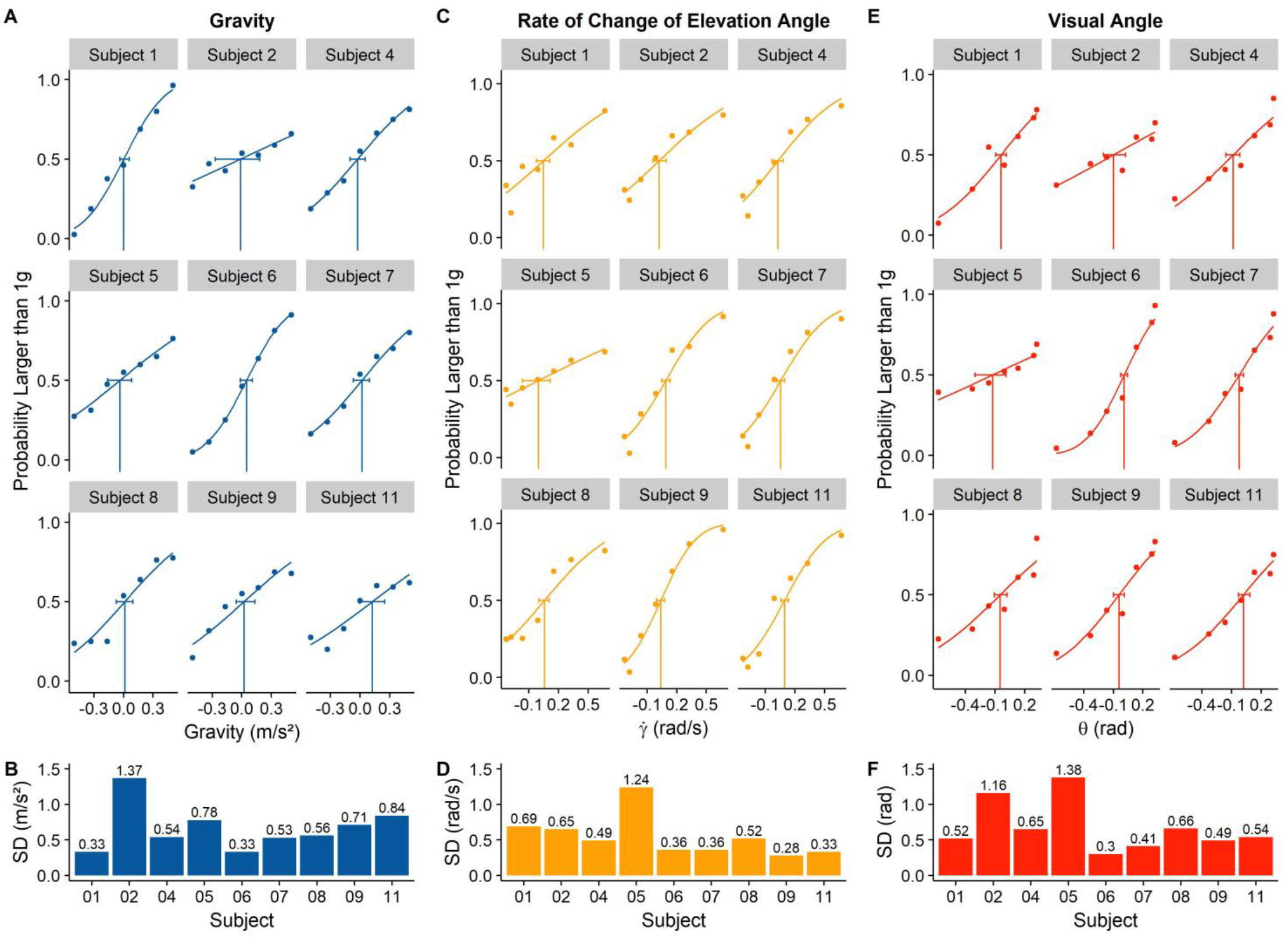
**A, C, E**. Psychometric functions based on gravity (A), difference in 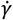 at 75 % of the trajectory (C) and difference in *θ* at 75 % (E) as decision variable for all participants included into analyses. The stimulus strength is plotted against the probability to judge the test parabola as having the higher underlying gravity. The horizontal bars indicate the confidence intervals for the PSEs. **B, D, F**. Standard deviations of the respective psychometric functions. All stimulus values were standardized by dividing them by the stimulus range to make SDs comparable across decision variables.

e then further analyzed the chosen models (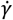 at 75 % and *θ* at 75 %) for differences in cue reliance with regards to initial speeds (see Table 2). 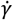 had a better model fit for *ν*_*ν*0_ = 3.7 m/s than for *ν*_*ν*0_ = 5.2 m/s. This trend was largely confirmed on the participant level. Regarding the two horizontal velocities, fits were superior for *ν_h_* = 6 m/s in comparison to *ν_h_* = 8.33 m/s. Here, some heterogeneity could be observed on the participant level: AIC differences (*AlC_ν_h_=8.33 m/s_* − *AlC_ν_h_=6.00 m/s_*) ranged between −10.7 for participant s01 and +11.6 for participant s02. For *θ*, model fits were better for *ν*_*ν*0_ = 5.2 m/s than for *ν*_*ν*0_ = 3.7 m/s and better for *ν_h_* = 8.33 m/s than for *v_h_* = 6 m/s. On the participant level, both trends proved relatively homogeneous.

**Table 2:** Breakdown of group AICs per initial vertical and horizontal velocities and decision variables.

| | $v_{v0} = 3.7 \text{ m/s}$ | $v_{v0} = 5.2 \text{ m/s}$ | $v_h = 6 \text{ m/s}$ | $v_h = 8.3 \text{ m/s}$ |
| --- | --- | --- | --- | --- |
| $\dot{\gamma}$ | **67.8** | 96.4 | **67.1** | 110.6 |
| $\theta$ | 94.3 | **68.4** | 96.2 | **72.1** |

Overall, the optical cue analyses allow for several conclusions. Firstly, information from later parts of the trajectory has a privileged role in gravity judgements. We thus picked 75 % of the trajectory as point of interest for further analyses. A comparison on the subject-level suggested that 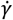 accounted for performance better than the placeholder decision variable gravity. However, other subjects’ performance was better explained by gravity, which is also supported by generally superior model fits for gravity with regards to 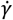 or *θ*, indicating that at least some subjects combine these cues to make more accurate judgements. Finally, fitting the data split by initial velocities suggests that subjects relied more on *γ* for lower and shorter parabolas (lower initial vertical and horizontal speeds) and more on *θ* for higher and longer parabolas (higher initial vertical and horizontal speeds).

#### A comparison of different cues to approximate an accurate gravity estimate

We then proceeded to further exploratory analyses of simple combinations of optic flow cues that may be good indicators of the gravity of a parabola, while avoiding the complexity of Equations (1) through (4). We are starting from the GS model (Gómez & López-Moliner, 2013), a model that has been used to describe how participants can extract TTC for parabolic trajectories from optical cues, assuming they have knowledge about gravity and the objects size.

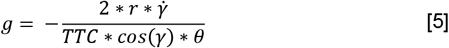

When solving the TTC model for gravity (Equation 5), it makes useful predictions about the underlying gravity value (see Figure 5). The model is tailored for parabolas that start and finish on the observer’s eye-height; a generalization of the model would thus be necessary for our stimulus, which originate and terminate below eye-height. But even in its less general form, it can provide an idea of how optical variables can be combined to obtain approximate estimates of gravity values: while the estimated values are generally higher than actual values and aren’t constant throughout the trajectories (see Figure 5), it still predicts accurately which of two parabolas has the higher underlying gravity. We thus fitted further psychometric functions for 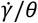 and a full GS model-based g estimate (“GS estimate”; Eq. 5). On the group level, 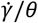 had a better model fit than 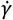 alone (AICs of 108.2 versus 126.4), while the GS estimate’s fit was superior to both (AIC of 78) and even approached the model fit of the placeholder decision variable Gravity (AIC of 58.7). On the subject level, the results were mixed (see Table 3), but the pattern was generally reproduced. This indicates that at least some subjects may be taking *γ* as base for their judgements and then rely on *θ* and/or TTC information compensate for ambiguities in *γ* information. When analyzing the SDs of the psychometric functions, the differences between 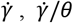 and GS estimate were generally neglectable. In comparison to Gravity, an interesting, but heterogeneous pattern emerged: The psychometric functions of some participants (s01, s05 and, to a lesser extent, s06) displayed lower SDs for gravity than for the three other, 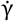-based decision variables. Other participants (like s09 and s11) had much lower SDs for 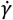-based decision variables. Interestingly, there was a (non-significant) tendency for participants with a higher overall performance, to have lower SDs for Gravity than for 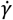-based decision variables (r = 0.44, p = 0.24 for 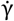; r = 0.49, p = 1.8 for 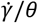; r = 0.39, p = 0.3 for GS estimate). Taken together, this may indicate that less successful participants relied too much on elevation angle information and failed to integrate it properly with the other available cues.

**Figure 5.**
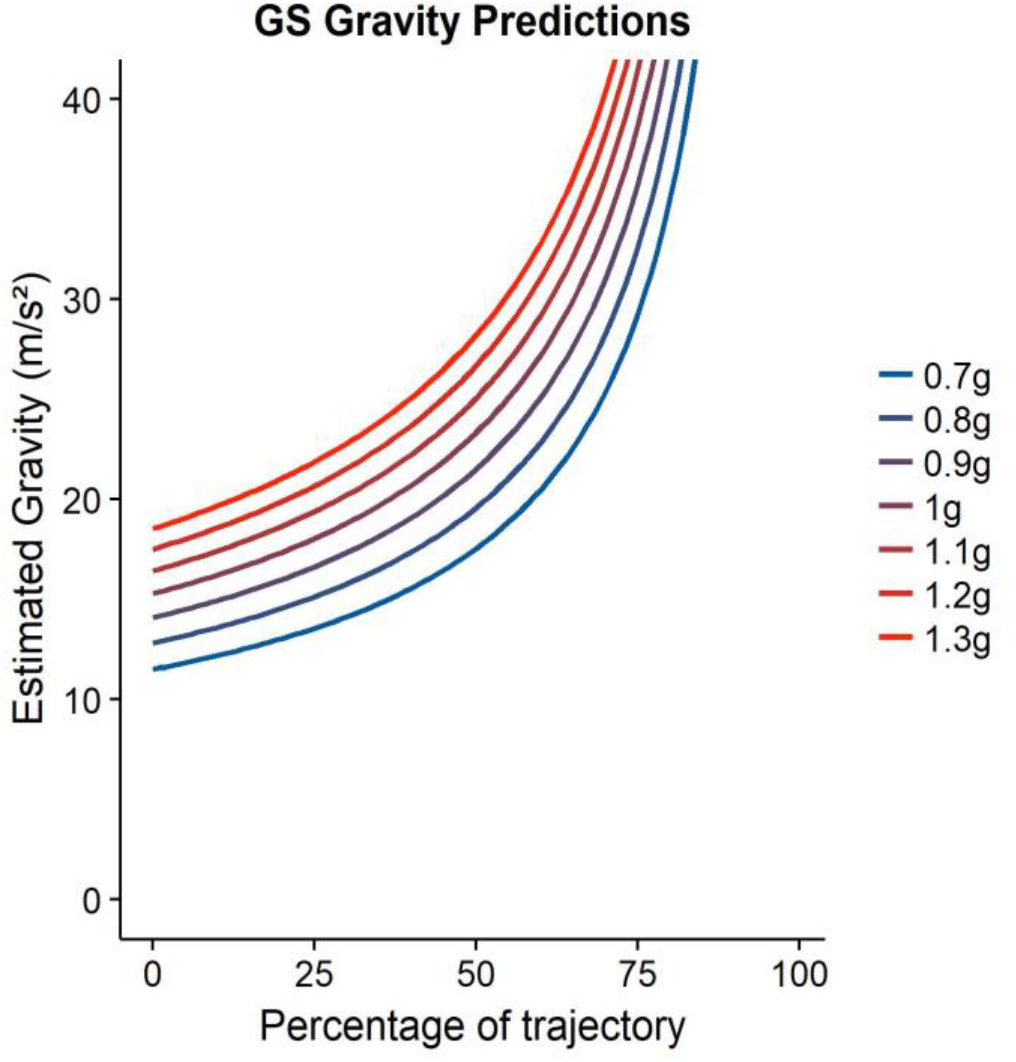
The lines indicate the predictions of the GS model for different gravities throughout the parabolas, *ν*_*ν*0_ = 5.2 m/s and *ν_h_* = 6 m/s. Patterns differed only minimally for the other velocity profiles.

**Table 3:**
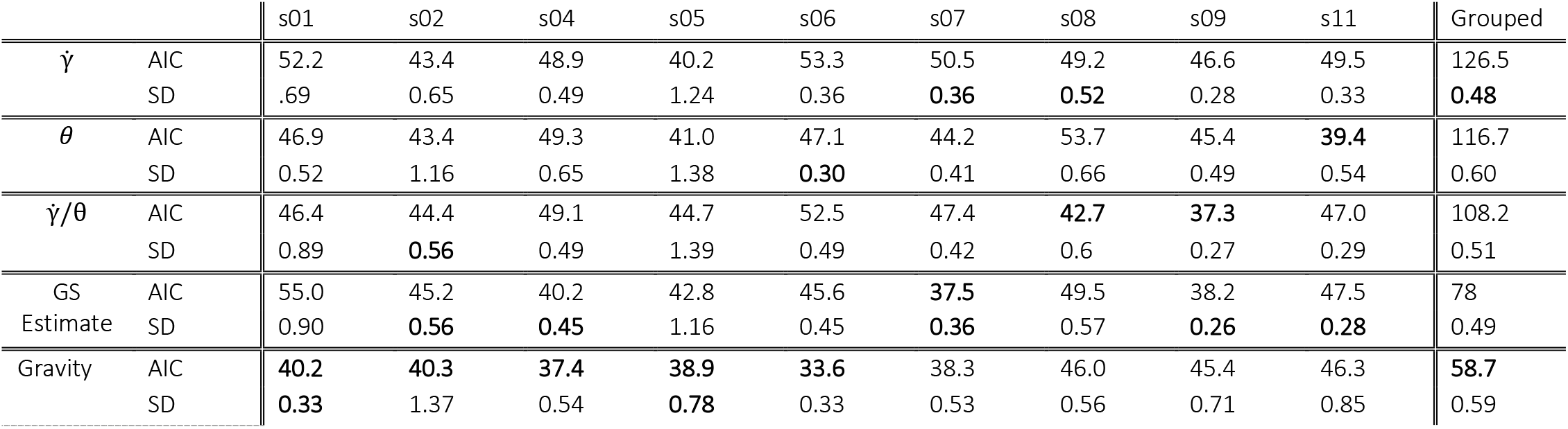
AICs and SDs for psychometric functions based on different decision variables at 75 % of the trajectory, both for individual subject and the all subjects grouped together. Lowest AIC and SD per subject are marked in bold.

#### Heterogeneity of strategy use

The previous results show there are some individual differences in the information used to judge gravity. In order to analyze these differences more systematically we employed General Linear Mixed Modelling using the MixedPsy package for R (see Moscatelli & Lacquaniti, 2012). We fitted a GLMM with subjects as grouping factor, and 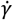 and *θ* both as fixed effects and as random effects, that is we allowed the intercept and slope?) to vary across the subjects. We compared this model with a GLMM in which there were not random variations of 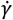 and *θ*, that is they were treated as fixed effects only. The GLMM with 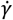 and *θ* as random effects had the superior model fit (AICs of 1760.9 versus 1796.2), and an ANOVA confirmed that it was significantly (p < 0.001) better than the comparison model.

#### Temporal information

As in our set of stimuli the total motion time correlated strongly with the gravity value (r = −0.76), subjects could have achieved acceptable performance levels by solely relying on this cue. To examine this possibility, we fitted a psychometric model across all subjects, using the overall motion time as decision variable. The goodness of fit of the model was worse (AIC of 210) than for all other decision variables (58.7 for gravity, 126.5 for 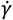 and 116.7 for *θ)*, which suggests that this cue played a subordinated role with regards to elevation angle and visual angle information.

Experiment 1 shows that elevation angle and visual angle information plays a role in gravity judgements. From previous models (e.g. the GS model mentioned above) and Equations 1 – 4, we know that also a correct representation of the target size can be of relevance. When size is known, the visual angle can be used to recover the remaining distance unambiguously, which in turn may aid the recovery of the underlying gravity. In Experiment 2, we thus manipulate size reliability.

## Experiment 2

Previous research (López-Moliner et al., 2007; López-Moliner & Keil, 2012) has shown that participants quickly assume a relatively constant ball size. We hypothesized therefore that participants in Experiment 1 maintained a representation of the ball size to judge gravity visually through the rate of change of the elevation angle and the rate of change of the visual angle (see also Equations 1 through 4). To determine the importance of a reliable and adequate representation of target size, we conducted a follow-up experiment with two different degrees of variability in target size.

### Materials and Methods

#### Participants

A total of twelve (n = 12) participants performed the task, among them two of the authors (BJ and LS). One of the authors (BJ) had also been a participant in Experiment 1 before. All participants had normal or corrected-to-normal vision. Two participants were excluded from the final analysis because they displayed chance level performance across all tested stimulus strengths. The remaining participants were between 21 and 30 years old and six (n = 6) were female. All participants gave their informed consent. The research in this study is part of an ongoing research program that has been approved by the local ethics committee of the University of Barcelona. The experiment was conducted in accordance with the Code of Ethics of the World Medical Association (Declaration of Helsinki).

#### Apparatus

The same setup was used as in Experiment 1.

#### Stimuli

The stimuli were identical to those from Experiment 1, with the difference that the target size of the test parabolas could be drawn either from a Gaussian with a mean of 0.033 m and a SD of 0.2 m (high variability condition, “HV”) or from a Gaussian with a mean of 0.033 m and a SD of 0.05 m (low variability condition, “LV”). Note that the size for the standard parabola remained fixed to 0.033 m. All trials containing a target radius below 0.004 m (103 trials across all participants, which corresponds roughly to 0.9 % of all trials) were excluded from the analysis because the targets were hardly visible.

#### Procedure

Participants performed first four blocks in the LV condition, with a total of 560 trials, and then four blocks in the HV condition, also with a total of 560 trials, or vice-versa. The order of presentation was counter-balanced such that five participants started with the LV condition and five started with the HV condition. Apart from these changes, the procedure was identical to Experiment 1.

### Results

As in Experiment 1, we employed the R package QuickPsy (Linares & López-Moliner, 2016) to fit psychometric functions to our data. In order to assess differences in discrimination thresholds and biases for target size variability and target size in gravity judgments, we compared low and high variability and small and big targets (see Figure 6). As for Experiment 1, we used the Bootstrap (Efron & Tibshirani, 1998) implemented in QuickPsy to compare PSEs and differences in discrimination thresholds between the respective conditions.

**Figure 6:**
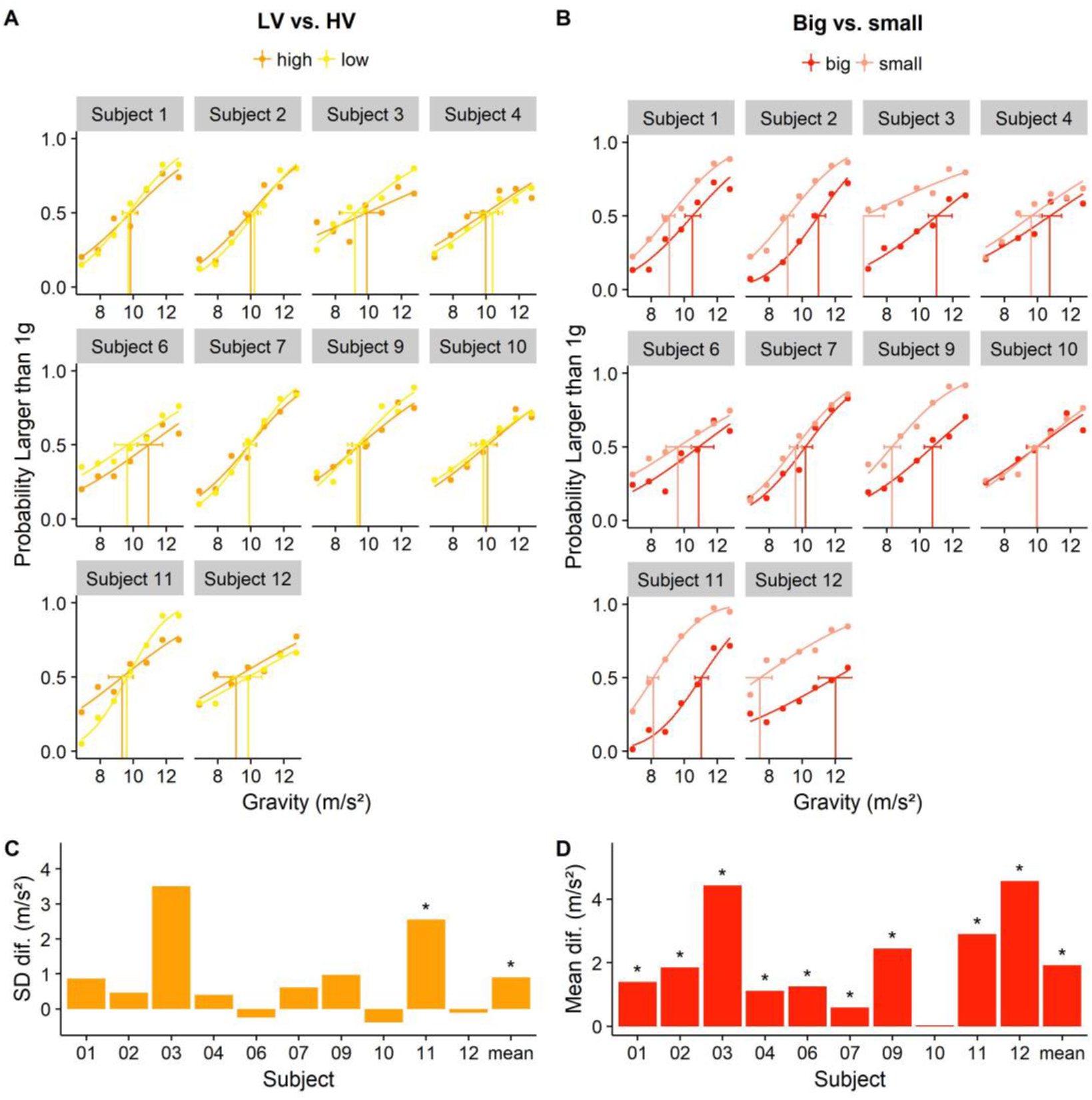
**A**. Psychometric functions per subject for Low variability (LV) vs. high variability (HV) of the ball size. **B**. Psychometric functions per subject for Big target size (> mean target size) vs. small target size (≤ mean target size). For both A. and B., the stimulus strength is plotted against the probability to judge the test parabola as having the higher underlying gravity. Horizontal bars indicate the confidence intervals around the respective PSEs. **C**. SD differences between LV and HV conditions per subject and for the whole group. **D**. SD differences between trials with Big and Small targets per subject and for the whole group.

Firstly, we evaluated the impact of target size variability, comparing low and high variability conditions for each subject (see Figure 6A/6C). Overall, subjects performed significantly better in the Low Variability condition (positive SD differences in Figure 6C). In individual analysis, after correcting for multiple comparisons by using a significance level of 1-0.05/n for the confidence intervals (with n = 10 comparisons, for a significance level of p = 0.995), only one subject (s11) showed a significant discrimination advantage for the LV condition. PSEs did not differ significantly.

Secondly, we assessed the impact of the ball size on the judgements of gravity (see Figure 6B/6D), comparing big vs. small balls. In the light of the GS model (Equation 5), if participants assume the same physical size, the variations of theta would induce a greater proportion of larger gravity responses for small sizes (which will be interpreted as the same size but underlying smaller visual angles). We established size categories with the mean ball size as cut-off criterion: “big” was defined as bigger than the mean ball size and “small” was defined as being less or equal the mean ball size. On the group-level, a significant bias was observed to judge small balls as having the bigger underlying gravity, as predicted by using the GS model while keeping size constant. Individual analyses confirmed, after correcting for multiple comparisons (see above, significance level of p = 0.995), that all participants except s10 displayed this bias. We did not find differences in discrimination thresholds between the two categories.

Finally, we compared the goodness of fit of psychometric models with 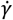 and *θ* as decision variables for big and small targets. The 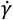 based model had a better fit for “Small” (AIC of 117.4) than for “Big” (AIC of 147.1), indicating that participants relied more on elevation angle information when the target was smaller. The *θ* model, in turn, had a better fit for “big” (AIC of 76.9) than for “small” (AIC of 131.2), indicating that participants relied more on visual angle information when the targets were bigger.

## Discussion

### High discrimination thresholds in visual gravity judgments

It is important to add that the mean PSE of 9.91 m/s^2^ (indicating high precision in the judgments) observed in Experiment 1 can not be interpreted meaningfully: the PSE indicates the stimulus strength at which it is equally likely for participants to answer that the test parabola had the higher underlying gravity as it is to answer that the standard parabola had the higher underlying gravity. A significant deviation from the mean (9.81 m/s^2^) would thus mean that participants were, for some reason beyond the stimulus strength alone, overall more likely to choose standard or test parabola. As the stimulus strength, at least in this analysis, is the only dimension that differs between test and standard parabola (all other parameters, such as initial velocities, are counter-balanced across trials), a PSE very close to the actual value of 9.81 m/s^2^ was expected.

While the discrimination thresholds for gravity judgments are in line with those observed for linear accelerations, the complexity of the underlying computations deserve some additional attention: even when an accurate representation of size is maintained, as we assume for the present experiment, two variables, the elevation angle, the visual angle, must be estimated and combined adequately. For an accurate estimate of gravity, the visual system would have to go through a complex series of computations: it needs to recover the position of the object in depth, its physical vertical velocity and finally it needs to compute the temporal derivative of the vertical speed. Prima facie, there are no significant obstacles for the visual system in the estimation of elevation angle and its derivative: the elevation angle can be recovered easily from first order information about the objects position with regards to the observer and its derivative corresponds to the retinal velocity in vertical direction. Indeed, 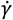 is, depending on the velocity profile, well above threshold for the last 50 % of the trajectory (McKee, 1981). On the other hand, while we pick up the visual angle itself readily, its temporal derivative is notoriously unreliable until the object gets very close to the observer (Regan & Hamstra, 1993). Note however that unlike in catching tasks, where neural delays render online visual information useless during the last 100 – 150 ms (Carlton, 1981) and where participants may not follow the target successfully with their gaze (Cesqui, Mezzetti, Lacquaniti, & D’Avella, 2015), late information may be exploited for the task at hand. Elevation Angle and Visual Angle information are therefore, in principle, available to the visual system with a good or at least a sufficient signal-to-noise ratio. It is therefore more likely that uncertainty arises from the correct integration of the available cues; the huge inter-subject variability when it comes to discrimination thresholds and cue reliance lend this claim additional support. A caveat in this respect is, certainly, that we did not record the participants eye movements. Our analysis is thus based on the assumption that they followed the target with their gaze throughout the trajectory, enabling them to collect the optic information.

### Decision variables and integration

As to the relevant cues for gravity judgements, an a priori analysis of how the stimuli unfold in space shows that both elevation angle (*γ*) and the visual angle (*θ*) information can contribute to extracting the underlying gravity of parabolic motion (see Figure 2 and Equations 1 – 4). Our data indicate that among elevation angle information, the temporal derivative 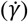 plays a prominent role, and among visual angle information, no clear prevalence of *θ* or 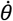 could be established. Theory, namely its use in the GS model for TTC estimation brought forward in (Gómez & López-Moliner, 2013), suggests, however, that *θ* may be privileged among visual angle-based cues. Also the motion duration was taken into account, but it was clearly less important than 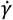 and *θ*. Since, however, the psychophysical models based on gravity as (placeholder) decision variable had superior model fits with regards to 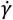 or *θ*, it is evident that none of the optic variables on their own can account for participants’ performance. While 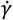 may account for the whole performance of some subjects, participants possibly use 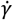 as principal cue and then add information from *θ* to achieve more accurate gravity estimates. This primacy of 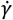 is supported by the observation that small target sizes in Experiment 2, which should make the estimation of the visual angle more noisy, did not lead to higher discrimination thresholds with regards to big target sizes. However, as this was not the main objective of Experiment 2, possible confounds (such as perceived distance) may cast some doubt on this conclusion.

We should reiterate, at this point, that we did not measure eye-movements. Therefore, our optic cue analyses are based on the assumption that participants followed the target with sufficient accuracy and precision. We furthermore assume that participants estimate the elevation angle using the starting position of the ball as reference, that is, we define the elevation angle as the angle between the line given by the observer’s eyes and the starting point on the parabola on the one hand, and the line given by the observer’s eyes and the target on the other hand. We therefore work under the hypothesis that the objective visual cues on which we based our analyses were sufficiently similar to the subject visual cues perceived by the participants. While we have no direct evidence to support these assumptions, we do believe that the relatively good model fits for the optic parameters lend this claim some additional credibility.

Incoherent discrimination threshold patterns for the psychometric functions fitted for gravity, 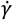 and *θ* between subjects indicate that they did not follow a unified strategy regarding the integration of cues, but rather combined them idiosyncratically. A GLMM analysis confirmed this tendency. Additionally, we could determine that there are not only inter-subject differences in cue reliance, but that also the characteristics of the parabola have an impact: elevation angle information was more important for lower vertical and lower horizontal velocities, that is, for lower and spatially shorter parabolas. This seems rather unintuitive, as steeper parabolas (that is higher initial vertical and lower horizontal velocities), should prima facie favor the recovery of 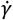. However, as evident from Figure 2, different gravities made a slightly bigger impact on 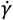 for lower initial vertical and horizontal velocities, which may explain this seeming anomaly. Conversely, participants relied more on *θ* for higher vertical and higher horizontal velocities, that is, for higher and spatially longer parabolas.

We furthermore demonstrate in Experiment 2 that an accurate representation of the ball size has a moderate influence on human discrimination thresholds in visual gravity judgments. Lower discrimination thresholds for small variability in ball size in comparison to high variability in ball size can be attributed to the brain maintaining one average representation of all presented ball sizes (López-Moliner & Keil, 2012). With a stable perceived physical ball size, differences in visual angle lead the brain to place the targets closer or further away in space, respectively. The same retinal speeds are therefore interpreted as higher physical speeds when the target is further away, which in turn gives rise to biases in gravity judgments: smaller balls are consistently judged as being governed by the higher underlying gravity and vice-versa. Experiment 2 also provided evidence that, when the targets were smaller, participants relied more on elevation angle information and less on visual angle information. This is expected, as observed visual angles are smaller and therefore less reliable for smaller targets, and as one source of information grows more unreliable, participants switch to other available cues, which is in our task mainly the elevation angle.

### Timing of information sampling

As mentioned above, we chose the optic variables at 75 % of the trajectory to fit the psychometric functions. We based this choice on a series of both a priori and a posteriori considerations. First of all, a geometric analysis of our stimulus trajectories (see Fig. 2) revealed that differences between the gravity levels both in elevation angle and visual angle information became more pronounced over the course of the trajectories. Furthermore, the signal-to-noise ratio of each optic variable grows as the target moves along its trajectory: depending on the velocity profiles, changes in *γ* are below discrimination threshold, as reported in (McKee, 1981), for some time within 25 % – 50 % of the trajectory, and then after about 60 % – 70 % its estimate becomes more noisy (de Bruyn & Orban, 1988). *θ* could generally be perceived with sufficient accuracy (McKee & Welch, 1992), while bigger absolute values later in the trajectory should facilitate making out differences. Furthermore, as shown in the results section of Experiment 1, models based on optic values further along the trajectory generally had better fits than those for other parts of the trajectory. Nonetheless, two caveats are in order: Firstly, the optic variables at 75 % correlate more strongly with gravity than at 60 %. Since gravity as placeholder decision variable had by far the best overall model fit, strong correlations of other decision variables with gravity could improve their model fits unduly. Secondly, Cesqui et al. (2015) observed in a catching experiment that smooth pursuit breaks down in the last part of parabolic trajectories. During this break-down phase, subjects generally perform a catch-up saccade (between 0.5 and 0.2 s before time-to-contact) and then fail to foveate the target altogether in a non-tracking period (0.15 s before time-to-contact until time-to-contact). Thus, for our stimuli, whose motion duration ranged between 0.58 and 1.5 s, the chosen region of interest (75 % of the trajectory) may fall into the catch-up saccade; nonetheless, subjects could still sample information before and after their catch-up saccade, thus taking advantage of the part of the trajectory that is richest in information. Furthermore, their experiment substantially differs from ours in that their participants performed interceptive movements, while our task was purely perceptual. Participants of Cesqui et al. (2015) may have not pursued the target in the last 0.15 s because neural delays rendered further information useless for catching at this point (Carlton, 1981). Taken together, there is thus sufficient evidence to support the claim that later information has a privileged role in the decision processes recruited in our task.

### Evidence for a weak likelihood

A more theoretical motivation for the present study was to determine qualitatively the reliability of the Bayesian likelihood for gravity perception. In this respect, it is important to point out that we are dealing with a highly complex “compound likelihood”: On the one hand, the brain integrates visual information from different sources such as the elevation angle and the visual angle. On the other hand, even for purely visual tasks such as the present experiment, cues from other modalities such as vestibular or bodily information impact perception (f. e. Senot et al., 2012; for a concise overview see also Lacquaniti et al., 2013). Furthermore, the overall reliability of the “gravity likelihood” aggregates not only uncertainty from perceiving these basic building blocks of information, but also from the integrative-decisional mechanism that combines all available basic information into a final judgment.

While Weber fractions of 13% to beyond 30% and the high inter-subject variability indicate that the overall precision of the visual system is relatively low for gravities, it is important to keep in mind that the psychophysically measured performance is a snapshot of the posterior reliability, which in turn is based on the reliability of likelihood and prior. The higher the reliability of likelihood and/or prior, the higher the reliability of the posterior. The evidence reviewed in (Jörges & López-Moliner, 2017) indicates that prior expectations about gravity have an important impact on posterior percepts of visually perceived gravity, which suggests that the prior attracts the posterior much more strongly than the likelihood (“strong prior”). Qualitatively, there are thus three possible scenarios: (1) Weak likelihood and medium prior, (2) weak likelihood and strong prior and (3) medium likelihood and strong prior. Of these tree possibilities, (3) is least compatible with the data collected in our experiments: while a medium likelihood and a strong prior would lead to a highly reliable posterior, our data suggest that the posterior is relatively unreliable, as evidenced by low precision and high inter-subject variability. Regarding (1) and (2), further experimental work will be necessary, as the present data does not allow to dissociate both possibilities.

## Conclusions

Humans have relatively high discrimination thresholds for the visual discrimination of different gravities expressed in parabolic motion. The two main sources of information available from optic flow, the rate of change of the Elevation Angle and the Visual Angle, were both important cues; our results suggest a primacy of the rate of change of the Elevation Angle, while the Visual Angle may be used to remedy its ambiguity with regards to underlying gravity values. We found evidence that the later parts of the stimulus trajectories represent a privileged source of information. As humans are sufficiently sensitive to all sources of information necessary to extract gravity for the visual scene, the most important source for variability is how the different primary cues are integrated. Translated into a Bayesian framework, our results could provide some evidence for a weak likelihood.

Beyond the theoretical implications described above, the present results represent a caveat for developers of virtual or augmented reality applications. They should expect users to face significant difficulties in environments with earth-discrepant visual gravities. While previous research (Zago et al., 2004; Zago & Lacquaniti, 2005) has shown that a full adaptation to 0 g might never occur while bodily gravity sensors signal 1 g, further research into the adaptation dynamics of immersive virtual environments might be useful to determine to what extent it is feasible to employ earth-discrepant gravities in such applications.

### Author Contributions and Notes

BJ conducted and analyzed Experiment 1 and wrote the paper. LS and BJ conducted and analyzed Experiment 2 in a joint effort. JLM provided the initial research question, programmed the stimuli and provided overall advice.

The authors declare no conflict of interest.

## Acknowledgments

Funding was provided by the Catalan government (2017SGR-48) and Ministry of Economy and Competition of the Spanish government: PSI2017-83493-R. The first author (BJ) was supported by an FI fellowship (FI-DGR 2016) from the Catalan government.

## References

Baurès, R., Benguigui, N., Amorim, M. A., & Siegler, I. A. (2007). Intercepting free falling objects: Better use Occam’s razor than internalize Newton’s law. Vision Research, 47, 2982–2991. https://doi.org/10.1016/j.visres.2007.07.024

Benguigui, N., Ripoll, H., & Broderick, M. P. (2003). Time-to-contact estimation of accelerated stimuli is based on first-order information. Journal of Experimental Psychology. Human Perception and Performance, 29(6), 1083–1101. https://doi.org/10.1037/0096-1523.29.6.1083

Bland, J. M., & Altman, D. G. (1995). Multiple significance tests: The Bonferroni method. Bmj, 310(6973), 170. https://doi.org/10.1136/bmj.310.6973.170

Brenner, E., Rodriguez, I. A., Muñoz, V. E., Schootemeijer, S., Mahieu, Y., Veerkamp, K., … Smeets, J. B. J. (2016). How can people be so good at intercepting accelerating objects if they are so poor at visually judging acceleration? I-Perception, 7(1), 1–13. https://doi.org/10.1177/2041669515624317

Brouwer, A.-M., Brenner, E., & Smeets, J. B. J. (2002). Perception of acceleration with short presentation times: can acceleration be used in interception? Perception & Psychophysics, 64(7), 1160–1168. https://doi.org/10.3758/BF03194764

Carlton, L. G. (1981). Processing visual feedback information for movement control. Journal of Experimental Psychology: Human Perception & Performance, 7, 1019–1030.

Cesqui, B., Mezzetti, M., Lacquaniti, F., & D’Avella, A. (2015). Gaze behavior in one-handed catching and its relation with interceptive performance: What the eyes can’t tell. PLoS ONE, 10(3), 1–39. https://doi.org/10.1371/journal.pone.0119445

de Bruyn, B., & Orban, G. A. (1988). Human velocity and direction discrimination measured with random dot patterns. Vision Research, 28(12), 1323–1335. https://doi.org/10.1016/0042-6989(88)90064-8

De Sá Teixeira, N. A., Hecht, H., & Oliveira, A. M. (2013). The representational dynamics of remembered projectile locations. Journal of Experimental Psychology: Human Perception and Performance, 39(6), 1690–1699. https://doi.org/10.1037/a0031777

Efron, B., & Tibshirani, R. (1998). Introduction to the Bootstrap World. CRC Press. https://doi.org/10.1214/ss/1063994971

Gómez, J., & López-Moliner, J. (2013). Synergies between optical and physical variables in intercepting parabolic targets. Frontiers in Behavioral Neuroscience, 7(May), 46. https://doi.org/10.3389/fnbeh.2013.00046

Hosking, S. G., & Crassini, B. (2010). The effects of familiar size and object trajectories on time-to-contact judgements. Experimental Brain Research, 203(3), 541–552. https://doi.org/10.1007/s00221-010-2258-7

Indovina, I., Maffei, V., Bosco, G., Zago, M., Macaluso, E., & Lacquaniti, F. (2005). Representation of visual gravitational motion in the human vestibular cortex. Science (New York, N.Y.), 308(April), 416–419. https://doi.org/10.1126/science.1107961

Jörges, B., & López-Moliner, J. (2017). Gravity as a Strong Prior: Implications for Perception and Action. Frontiers in Human Neuroscience, 11(203). https://doi.org/10.3389/fnhum.2017.00203

Lacquaniti, F., Bosco, G., Indovina, I., La Scaleia, B., Maffei, V., Moscatelli, A., & Zago, M. (2013). Visual gravitational motion and the vestibular system in humans. Frontiers in Integrative Neuroscience, 7(December), 101. https://doi.org/10.3389/fnint.2013.00101

Lee, D. N., & Reddish, P. E. (1981). Plummeting gannets: a paradigm of ecological optics. Nature, 293, 293–294. https://doi.org/10.1038/293293a0

Linares, D., & López-Moliner, J. (2016). quickpsy: An R Package to Fit Psychometric Functions for Multiple Groups. The R Journal, 8(1), 122–131. Retrieved from https://journal.r-project.org/archive/2016-1/linares-na.pdf

López-Moliner, J., Field, D. T., & Wann, J. P. (2007). Interceptive timing: prior knowledge matters. Journal of Vision, 7, 1–8. https://doi.org/10.1167/7.13.11

López-Moliner, J., & Keil, M. (2012). People Favour Imperfect Catching by Assuming a Stable World. Current Science, (4), 1435–1439. https://doi.org/10.1371/Citation

López-Moliner, J., Maiche, A., & Estaún, S. (2003). Perception of acceleration in motion-in-depth with only monocular and both monocular and binocular information. Psicológica, 24, 93–108.

Maffei, V., Mazzarella, E., Piras, F., Spalletta, G., Caltagirone, C., Lacquaniti, F., & Daprati, E. (2016). Processing of visual gravitational motion in the peri-sylvian cortex: evidence from brain-damaged patients. Cortex, 78, 55–69. https://doi.org/10.1016/j.cortex.2016.02.004

McIntyre, J., Zago, M., & Berthoz, A. (2001). Does the Brain Model Newton’s Laws. Nature Neuroscience, 12(17), 109–110. https://doi.org/10.1097/00001756-200112040-00004

McIntyre, J., Zago, M., Berthoz, A., & Lacquaniti, F. (2003). The Brain as a Predictor: On Catching Flying Balls in Zero-G. In J. C. Buckey & J. L. Homick (Eds.), The Neurolab Spacelab Mission: Neuroscience Research in Space (pp. 55–61). National Aeronautics and Space Administration, Lyndon B. Johnson Space Center.

McKee, S. P. (1981). A local mechanism for differential velocity detection. Vision Research, 21, 491–500.

McKee, S. P., & Welch, L. (1992). The precision of size constancy. Vision Research, 32(8), 1447–1460. https://doi.org/10.1016/0042-6989(92)90201-S

Moscatelli, A., & Lacquaniti, F. (2012). Modeling psychophysical data at the population-level: The generalized linear mixed model, 12(2012), 1–17. https://doi.org/10.1167/12.11.26.Introduction

R Core Team. (2017). A Language and Environment for Statistical Computing. R Foundation for Statistical Computing,. Vienna, Austria. Retrieved from http://www.r-project.org/.

Regan, D., & Beverley, K. I. (1979). Binocular and monocular stimuli for motion in depth: Changing-disparity and changing-size feed the same motion-in-depth stage. Vision Research, 19(12), 1331–1342. https://doi.org/10.1016/0042-6989(79)90205-0

Regan, D., & Hamstra, S. J. (1993). Dissociation of discrimination thresholds for time to contact and for rate of angular expansion. Vision Research, 33(4), 447–462. https://doi.org/10.1016/0042-6989(93)90252-R

Senot, P., Zago, M., Le Seac’h, a., Zaoui, M., Berthoz, a., Lacquaniti, F., & McIntyre, J. (2012). When Up Is Down in 0g: How Gravity Sensing Affects the Timing of Interceptive Actions. Journal of Neuroscience, 32(6), 1969–1973. https://doi.org/10.1523/JNEUROSCI.3886-11.2012

Werkhoven, P., Snippe, H. P., & Alexander, T. (1992). Visual processing of optic acceleration. Vision Research, 32(12), 2313–2329. https://doi.org/10.1016/0042-6989(92)90095-Z

Zago, M., Bosco, G., Maffei, V., Iosa, M., Ivanenko, Y. P., & Lacquaniti, F. (2004). Fast Adaptation of the Internal Model of Gravity for Manual Interceptions: Evidence for Event-Dependent Learning. Journal of Neurophysiology, 93(2), 1055–1068. https://doi.org/10.1152/jn.00833.2004

Zago, M., & Lacquaniti, F. (2005). Internal Model of Gravity for Hand Interception: Parametric Adaptation to Zero-Gravity Visual Targets on Earth. Journal of Neurophysiology, 94(2), 1346–1357. https://doi.org/10.1152/jn.00215.2005

Zago, M., McIntyre, J., Senot, P., & Lacquaniti, F. (2008). Internal models and prediction of visual gravitational motion. Vision Research, 48(14), 1532–1538. https://doi.org/10.1016/j.visres.2008.04.005

